# Stacked Generalization: An Introduction to Super Learning

**DOI:** 10.1101/172395

**Authors:** Ashley I. Naimi, Laura B. Balzer

## Abstract

Stacked generalization is an ensemble method that allows researchers to combine several different prediction algorithms into one. Since its introduction in the early 1990s, the method has evolved several times into what is now known as “Super Learner”. Super Learner uses *V* -fold cross-validation to build the optimal weighted combination of predictions from a library of candidate algorithms. Optimality is defined by a user-specified objective function, such as minimizing mean squared error or maximizing the area under the receiver operating characteristic curve. Although relatively simple in nature, use of the Super Learner by epidemiologists has been hampered by limitations in understanding conceptual and technical details. We work step-by-step through two examples to illustrate concepts and address common concerns.

## KEY MESSAGES

- Stacked generalization is a meta-learning algorithm that allows researchers to combine several different methods to improve predictive performance.
- Since its introduction in the early 1990s, stacked generalization has evolved both practically and theoretically into the Super Learner.
- The Super Learner combines predictions from several methods by using *V* -fold cross-validation and minimizing a user-specified loss function, such as prediction error, negative-log-likelihood, or rank loss.
- Under reasonable constraints, Super Learner is guaranteed to perform asymptotically as well as the best performing algorithm included in the candidate set of algorithms.
- In practice, Super Learner is a powerful tool to estimate complex, real-world relationships while avoiding unsubstantiated parametric assumptions and over-fitting.

Predicting health-related outcomes is a topic of major interest in clinical and public health settings. Despite numerous advances in methodology in the past two decades, clinical and population health research scientists continue to rely heavily on parametric (e.g., logistic) regression models for prediction. Often, only a single model is specified to generate predictions.

In the early 1990s, Wolpert developed an approach to combine several “lower-level” machine learning methods into a “higher-level” model with the goal of increasing predictive accuracy. ^1^ He termed the approach “stacked generalization”, which later became known as “stacking”. Later, Breiman demonstrated how stacking can be used to improve the predictive accuracy in a regression context, and showed that imposing certain constraints on the higher-level model improved predictive performance. ^2^ More recently, van der Laan and colleagues proved that stacking possesses certain ideal theoretical properties. ^3–5^ In particular, their oracle inequality guarantees that in large samples the algorithm will perform at least as well as the best individual predictor included in the ensemble. Therefore, choosing a large library of diverse algorithms will enhance performance, and creating the best weighted combination of candidate algorithms will further improve performance. Here, “best” is defined in terms of a bounded loss function, and over-fitting is avoided with *V* -fold crossvalidation. In this context, the term “Super Learner” was coined.

Super Learner has tremendous potential for improving the quality of prediction algorithms in applied health sciences, and minimizing the extent to which empirical findings rely on parametric modeling assumptions. While the number of simulation studies and applications of Super Learner are growing, ^6–10^ wider use may be hampered by comparatively few pedagogic examples. Here, we add to prior work by providing step-by-step implementation in the context of two simple simulations. Our examples are motivated by common challenges to estimate a dose-response curve, and to build a classifier for a binary outcome. We also link Super Learner to prior work on stacking, provide guidelines on fitting the Super Learner to empirical data, and discuss common concerns. Full `R` code ^11^ is publicly available at GitHub.

## Example 1: Dose-Response Curve

For 1,000 observations, we generate a continuous exposure *X* by drawing from a uniform distribution with a minimum of zero and a maximum of eight and then generate a continuous outcome *Y* as

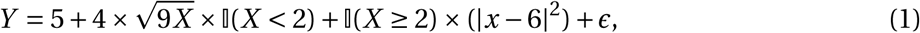

where []() denotes the indicator function evaluating to 1 if the argument is true (zero otherwise), and *£* was drawn from a doubly-exponential distribution with mean zero and scale parameter one. The true dose-response curve is depicted by the black line in Figure 1. We now manually demonstrate how Super Learner can be used to flexibly model this relation without making parametric assumptions.

**Figure 1:**
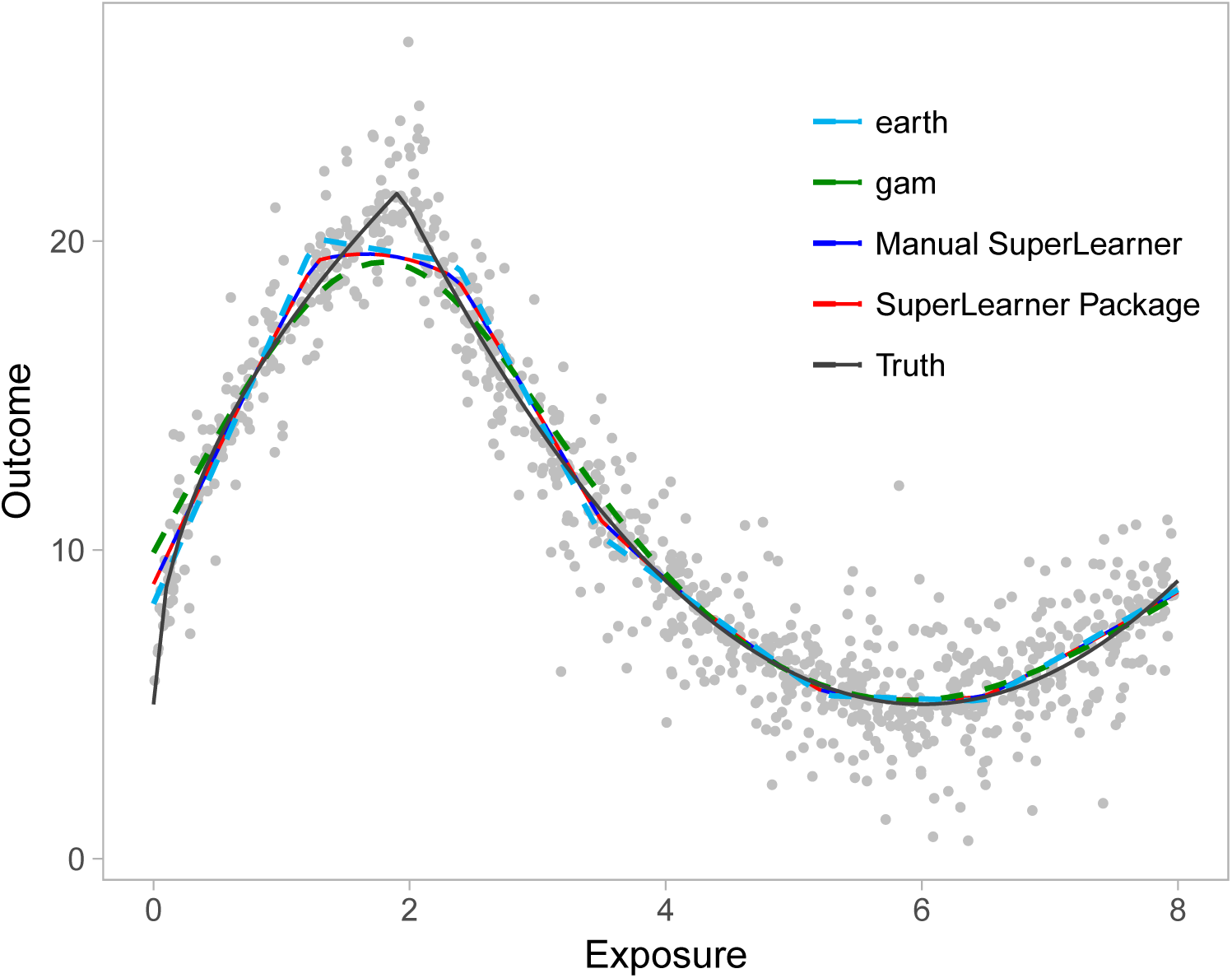
Dose-response curves for the relation between our simulated continuous exposure and continuous outcome in Example 1. The black line represents the true curve, while the red and blue lines represent curves estimated with the programmed Super Learner package in R, and the manually coded Super Learner. Light blue and green curves show the fits from the level-zero algorithms, `earth` and `gam` respectively. Gray dots represent observed data-points.

To simplify our illustration, we consider only two “level-zero” algorithms as candidates to the Super Learner library: generalized additive models with 5-knot natural cubic splines (`gam`)^12^ and multivariate adaptive regression splines implemented via the `earth` package. ^17^ In practice, a large and diverse library of candidate estimators is recommended. Our specific interest is in quantifying the mean of *Y* as a (flexible) function of *X*. To measure the performance of candidate algorithms and to construct the weighted combination of algorithms, we select the *L*-2 squared error loss function (*Y -Ŷ*)^2^ where *Ŷ* denotes our predictions. Minimizing the expectation of the *L*-2 loss function is equivalent to minimizing mean squared error, which is the same objective function used in ordinary least squares regression. ^13(section 2.4)^ To estimate this expected loss, called the “risk”, we use *V* -fold cross-validation with *V =* 5 folds.

**Step 1.** Split the observed “level-zero” data into 5 mutually exclusive and exhaustive groups of *n*/*V =* 1000/5 *=* 200 observations. These groups are called “folds”.

**Step 2.** For each fold *v =* {1,…, 5},

a. Define the observations in fold *v* as the validation set, and all remaining observations (80% of the data) as the training set.
b. Fit each algorithm on the training set.
c. For each algorithm, use its estimated fit to predict the outcome for each observation in the validation set. Recall the observations in the validation set are not used train each candidate algorithm.
d. For each algorithm, estimate the risk. For the *L*-2 loss, we average the squared difference between the outcome *Y* and its prediction *Ŷ* for all observations in the validation set *v*. In other words, we calculate the mean squared error (MSE) between the observed outcomes in the validation set and the predicted outcomes based on the algorithms fit on the training set.

**Step 3.** Average the estimated risks across the folds to obtain one measure of performance for each algorithm. In our simple example, the cross-validated estimates of the squared prediction error are 2.58 for `gam` and 2.48 for `earth`.

At this point, we could simply select the algorithm with smallest cross-validated risk estimate (here, `earth`). This approach is sometimes called the Discrete Super Learner. ^6^ Instead, we combine the cross-validated predictions, which are referred to as the “level-one” data, to improve performance and build the “level-one” learner.

**Step 4.** Let *Ŷ*_gam-cv_ and *Ŷ*_earth-cv_ denote the cross-validated predicted outcomes from `gam` and `earth`, respectively. Recall the observed outcome is denoted *Y*. To calculate the contribution of each candidate algorithm to the final Super Learner prediction, we use non-negative least squares to regress the actual outcome against the predictions, while suppressing the intercept and constraining the coefficients to be non-negative and sum to 1:

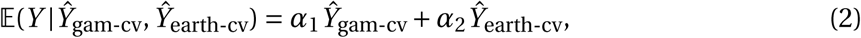

such that *α*_1_ *≥* 0; *α*_2_ *≥* 0, and 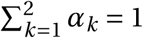&. Combining the Ŷ_gam-cv_ and Ŷ_earth-cv_ under these constraints (non-negative estimates that sum to 1) is referred to as a “convex combination,” and is motivated by both theoretical results and improved stability in practice. ^2,5^ Non-negative least squares corresponds to minimizing the mean squared error, which is our chosen loss function (and thus, fulfills our objective). We then normalize the coefficients from this regression to sum to 1. In our simple example, the normalized coefficient values are 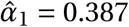 for `gam` and 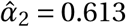 for `earth`. Therefore, both generalized additive models and regression splines each contribute approximately 40% and 60% of the weight in the optimal predictor.

**Step 5.** The final step is to use the above weights to generate the Super Learner, which can then be applied to new data (*X*) to predict the continuous outcome. To do so, re-fit `gam` and `earth` on the entire sample and denote the predicted outcomes as Ŷ_gam_ and Ŷ_earth_, respectively. Then combine these predictions with the estimated weights from Step 4:

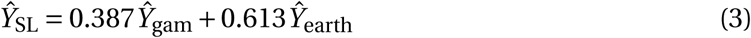

where Ŷ_SL_ denotes our final Super Learner predicted outcome. The resulting predictions are shown in blue in Figure 1. For comparison, the predictions from the `R` package `SuperLearner`v2.0-23-9000 are shown in red, ^14^ while the predictions from `gam` and `earth` are shown in light blue and green, respectively.

## Example 2: Binary Classification

Our second example is predicting the occurrence of a binary outcome with goal of maximizing the area under the receiver operating characteristic (ROC) curve, which shows the balance between sensitivity and specificity for varying discrimination thresholds. For 10,000 observations, we generate five covariates **X** *=* {*X*_1_,…, *X*_5_} by drawing from a multivariate normal distribution and then generate the outcome *Y* by drawing from a Bernoulli distribution with probability

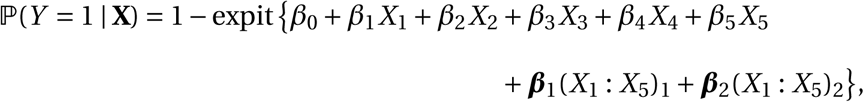

where expit(*•*) *=* (1 *+* exp[*-•*])^*-*1^; and (*X*_1_ : *X*_5_)_1_ and (*X*_1_ : *X*_5_)_2_ denote all first and second order interactions between *X*_1_,…, *X*_5_, respectively; and ***β***_1_ and ***β***_2_ denote a set of parameters, one for each interaction. In total, there were 25 terms plus the intercept in this model. The intercept was set to *β*_0_ *=* 2, while all other parameters were drawn from a uniform distribution bounded by 0 and 1.

Our library of candidate algorithms consists of Bayesian GLMs (`bayesglm)`, implemented via the `arm` package (v 1.9-3), ^15^ and multivariate polynomial adaptive regression splines (`polymars)` via the `polspline` package. ^16^ Our objective is to generate an algorithm that correctly classifies individuals given covariates. Correct classification is a function of both sensitivity and specificity. Thus, to measure the performance of these level-zero algorithms and build the meta-learner, we use the rank loss function, which corresponds to maximizing the area under the ROC curve (AUC). ^18^ We again use 5-fold cross-validation to obtain an honest measure of performance and avoid over-fitting.

**Steps 1-3.** The implementation of Steps 1-3 are analogous to the previous example. In place of `gam` and `earth`, we use `bayesglm`, and `polymars`. In place of the *L*-2 loss function, we use the rank loss. Specifically, for each validation set and each algorithm, we estimate the sensitivity, specificity, and then compute the AUC. The expected loss (i.e., “risk”) can then be computed as 1*-*AUC. Averaging the estimated risks across the folds yields one measure of performance for each algorithm. In our simple example, the cross-validated risk estimates are 0.122 for `bayesglm` and 0.114 for `polymars`.

As before, we could simply select the model with lowest cross-validated risk estimate (here, `polymars`). Instead in Steps 4-5, we combine the resulting cross-validated predictions to generate the level-one learner.

**Step 4.** Let Ŷ_bglm-cv_ and Ŷ_pm-cv_ denote the cross-validated predictions from `bayesglm` and `polymars`, respectively. To calculate the contribution of each candidate algorithm to the final Super Learner prediction, use the rank loss function to define “optimal” as the convex combination that maximizes the AUC. Then estimate the *α* parameters in the following constrained regression

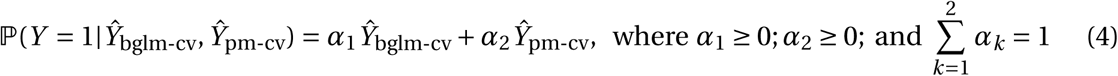

such that (1 *-AUC*) is minimized when comparing Super Learner predicted probabilities to the observed outcomes. These parameters can be estimated with a tailored optimization function, such as `optim`. In our simple example, the coefficient values are 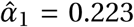 for `bayesglm` and 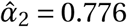 for `polymars`.

**Step 5.** As before, the final step is to use the above coefficients to generate the Super Learner. To do so, refit `bayesglm` and `polymars` on the entire sample and denote the predicted outcomes as *Ŷ*_bglm_ and *Ŷ*_pm_ Then combine these predictions with the estimated weights:

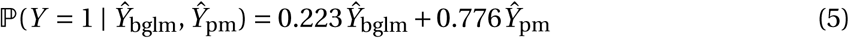

These predictions can then be used to compute the ROC curve displayed in blue in Figure 2. For comparison the predictions from the `R` package `SuperLearner` are shown in red.

**Figure 2:**
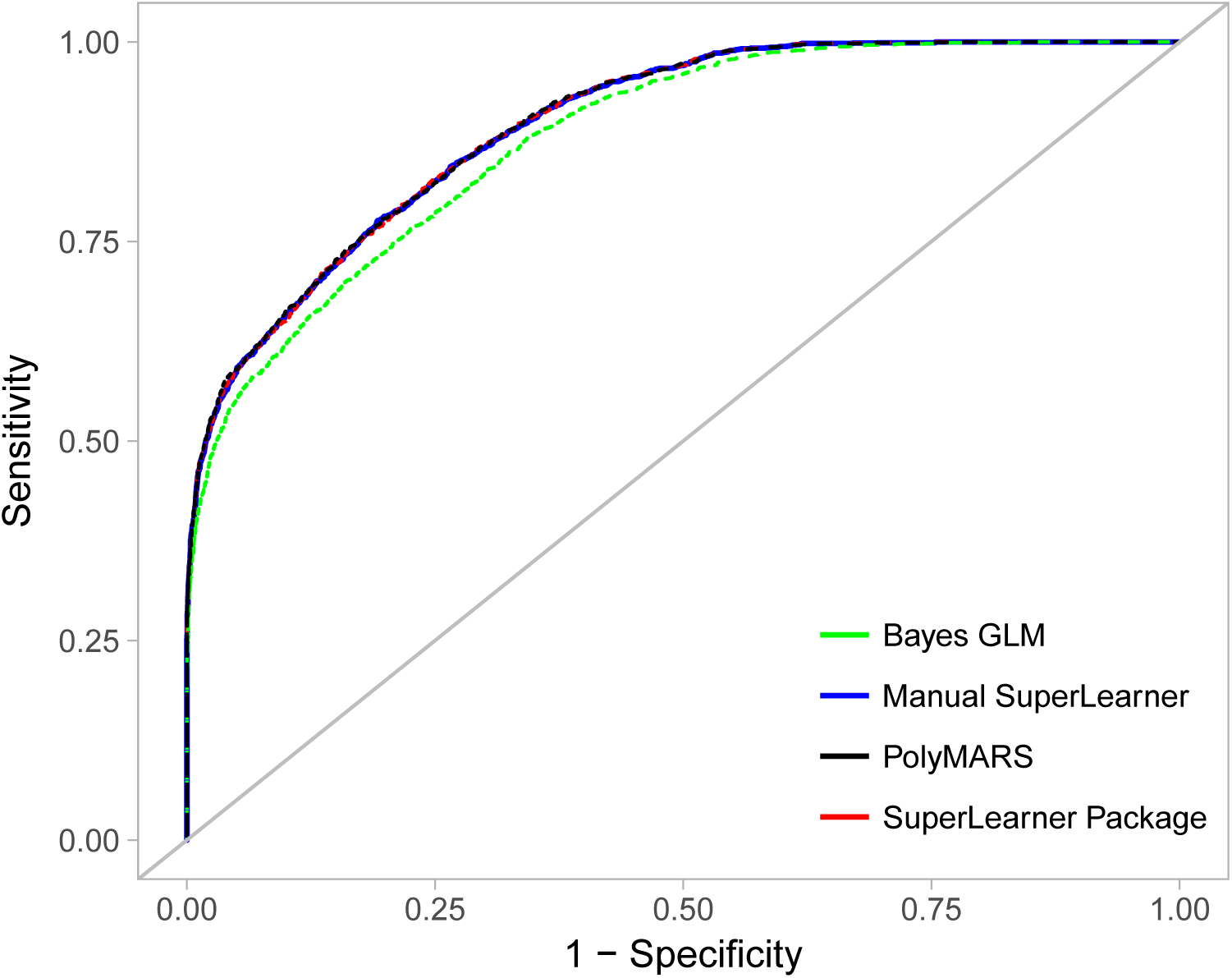
Receiver operating characteristic curves displaying the ability of 5 simulated exposures to predict the simulated outcome in Example 2. Blue line represents the curve obtained from the Super Learner package. Red dotted line represents curve obtained from manually coded Super Learner. The green line represents the curve from level-zero Bayes GLM algorithm, and the black line represents the curve from PolyMARS.

## Discussion

Stacked generalizations, notably the Super Learner, are fast becoming an important part of the epidemiologic toolkit. There are several challenges in understanding precisely what stacking is, and how it is implemented. These challenges include the use of complex machine learning algorithms as candidate algorithms, and the actual process by which the resulting predictions are combined into the meta-learner. Here, we sought to clarify the latter aspect, namely, implementation of the Super Learner algorithm.

Several considerations (including strengths and limitations) merit attention. First, any number of loss functions could be chosen determine the optimal combination of algorithm-specific predictions. The choice should be based on the objective of the analysis. The target parameter depends on the loss function choice, but various loss functions can identify the same target parameter as minimizer of its risk. Our examples demonstrated the use of the *L*-2 loss to minimize prediction error when estimating a dose-response curve, and the rank loss to maximize AUC when developing a binary classifier. Other loss functions could be entertained. For example, Zheng *et al.* recently aimed to simultaneously maximize sensitivity and minimize rate of false positive predictions with application to identify high-risk individuals for pre-exposure prophylaxis. ^10^

Second, a wide array of candidate algorithms can be included in the Super Learner library. We recommend including standard parametric modeling approaches (e.g., generalized linear models, simple mean, simple median) and more complex data-adaptive methods (e.g., penalized regression, and tree- or kernel-based methods). Often, the performance of data-adaptive methods depends on how tuning parameters are specified. The Super Learner can also include variations of the same algorithm under different tuning parameter choices (e.g., extreme gradient boosting with varying shrinkage rates). We refer the readers to Polley et al. ^14^ for a practical demonstration.

Third, a critically important part of Super Learning is the use of *V* -fold cross-validation. However, the optimal choice of *V* is not always clear. At one end of the extreme, leave-one-out cross validation chooses *V = N*, but is subject to potentially high variance and low bias. On the other end, 2-fold cross validation is subject to potentially low variance and high bias. A general rule of thumb is to use *V =* 10. ^19^ Though common, this number will not optimize performance in all settings. ^20^ In general, we recommend increasing the number of folds (*V*) as sample size *n* decreases.

While the Super Learner with a rich set of candidates represents a versatile tool, important limitations should be noted, particularly in the context of effect estimation. First, no algorithm (Super Learner or any other machine learning method) should be used to replace careful thinking of the underlying causal structure. The goal of Super Learner is to do the best (as specified through the loss function) possible job predicting the outcome (or exposure) given the inputted covariates. Super Learner does not distinguish between confounders, instrumental variables, mediators, and the exposure. Indeed, Super Learner is a“black box” algorithm; so the exact contribution of each covariate to prediction is unclear. These contributions can be revealed by estimating variable importance measures, which quantify the marginal association between each predictor and the outcome after adjusting for the others. Nevertheless, a large predictor contribution may be the result of direct causation, unmeasured confounding, collider stratification, reverse causation, or some other mechanism.

The goal of prediction is distinct from causal effect estimation, but prediction is often an intermediate step in estimating causal effects. ^31^ Indeed, some researchers have been advocating for the use of data-adaptive methods, including the Super Learner, for effect estimation via singly-robust methods, depending on estimation of either the conditional mean outcome or the propensity score. ^21–27^ While flexible algorithms can reduce the risk of bias due to regression model misspecification, a serious concern is that the use of data-adaptive algorithms in this context can result in invalid statistical inference (i.e. misleading confidence intervals). Specifically, there is no theory to support the resulting estimator is asymptotically linear (i.e., consistent and asymptotically normal). ^28,31^ This concern can be alleviated through the use of doubly robust estimators ^29–33^ or higher-order influence function based estimators. ^34,35^ In contrast, statistical inference is usually not included in prediction algorithms ^13^ (as in our examples). One could, however, evaluate the performance of Super Learner through an additional layer of cross-validation. ^6^

Overall, Super Learner is an important tool that researchers can use to improve predictive accuracy, avoid overfitting, and minimize parametric assumptions. We have provided a simple explanation of the Super Learner to facilitate a more widespread use in epidemiology. More advanced treatments with realistic data examples are available ^5,7,31^ and should be consulted for additional depth.

## Acknowledgements

We thank Susan Gruber andMark J van der Laan for expert advice.

## Sources of Funding

NIH grant number UL1TR001857 and R37AI051164

